# Genetic diversification of persistent *Mycobacterium abscessus* within Cystic Fibrosis patients

**DOI:** 10.1101/2021.02.19.431934

**Authors:** Astrid Lewin, Elisabeth Kamal, Torsten Semmler, Katja Winter, Sandra Kaiser, Hubert Schäfer, Lei Mao, Patience Eschenhagen, Claudia Grehn, Carsten Schwarz

## Abstract

*Mycobacterium (M.) abscessus* infections in Cystic Fibrosis (CF) patients cause a deterioration of lung function. Treatment of these multidrug-resistant pathogens is associated with severe side-effects, while frequently unsuccessful. Insight on *M. abscessus* genomic evolvement during chronic lung infection would be beneficial for improving treatment strategies. A longitudinal study enrolling 42 CF patients was performed at a CF center in Berlin, Germany, to elaborate phylogeny and genomic diversification of in-patient *M. abscessus*. Eleven of the 42 CF patients were infected with *M. abscessus*. Global human-transmissible *M. abscessus* cluster strains were isolated from five of these 11 patients. Phylogenetic analysis of 88 genomes from isolates of the 11 patients excluded occurrence of *M. abscessus* transmission among members of the study group. Genome sequencing and variant analysis of 30 isolates from 11 serial respiratory samples collected over four years from a chronically infected patient demonstrated accumulation of gene mutations. In total, 53 genes exhibiting non-synonymous variations were identified. Enrichment analysis emphasized genes involved in synthesis of glycopeptidolipids, genes from the *embABC* (arabinosyltransferase) operon, *betA* (glucose-methanol-choline oxidoreductase) and *choD* (cholesterol oxidase). Genetic diversity evolved in a variety of virulence- and resistance-associated genes. The strategy of *M. abscessus* populations in chronic lung infection is not clonal expansion of dominant variants, but to sustain simultaneously a wide range of genetic variants facilitating adaptation of the population to changing living conditions in the lung. Genomic diversification during chronic infection requires increased attention when new control strategies against *M. abscessus* infections are explored.

## Introduction

More than 30 species of nontuberculous mycobacteria (NTM) are known to cause infections in humans [1], mostly in persons with immunodeficiency or underlying diseases such as Cystic Fibrosis (CF) [2]. NTM prevalence in CF populations is increasing [3], which can in part be attributed to medical progress resulting in extended life expectancy of CF patients. Particularly, infections with *Mycobacterium abscessus* (MABS), also named *Mycobacteroides abscessus*, pose a threat to CF patients as they frequently lead to lung function decrease [4] and pose a risk in lung transplantation [5]. The environment has generally been assumed to be the source of NTM infection. However, recent findings show that the majority of MABS in CF patients belong to three phylogenetic clusters [6] suggesting human-to-human transmission as an additional potential infection route. MABS is characterized by its extreme resistance towards antibiotics [7], which necessitates long-lasting combination therapy. Despite such protracted therapy causing severe side effects, low culture conversion rates of typically 40-50% have been reported [8]. Because of the serious side effects of the antibiotic therapies, the balance between their harm and benefit for patients must be carefully weighed. Although information on genomic diversity of MABS is now available [6], prognostic genomic markers for disease progression have not yet been identified and knowledge on in-patient evolution of MABS during chronicity is scarce. Using respiratory samples from CF patients from a German CF treatment center, the focus of the present study lies on characterizing the population structure of MABS in CF patients and analyzing in-patient genetic variation evolving during long-term chronic infection.

## Materials and Methods

### Isolation and identification of NTM from Cystic Fibrosis patients

The 42 patients involved in the study were recruited from the CF Center at the Charité-Universitätsmedizin Berlin in Germany from 2013 to 2018. Within this period, the center treated 16 CF patients with NTM-PD. Patient characteristics are summarized in Supplementary Table S1. Isolation of NTM was envisaged if a patient had an unclear decline in pulmonary function tests that did not respond to bacteria-targeted antibiotics or during the annual check-up. Permission for the study was obtained from the ethics committee of the Charité –Universitätsmedizin Berlin (EA2/093/12). Written consent had been obtained from all patients.

Isolation of NTM from sputum or BAL using Nalc/NaOH was performed as described in [9]. NTM colonies isolated from sputum were purified at least twice by spreading single colonies on agar plates. NTM species were determined by PCR (DreamTaq DNA Polymerase, Thermo Fisher Scientific) and sequencing (ABI BigDyeTM 3.1, Thermo Fisher Scientific) of 16S rDNA and/or ITS. Primers used are listed in Supplementary Table S2.

### Cultivation of MABS

MABS was grown at 37°C on Middlebrook 7H11 agar (BD Biosciences) supplemented with 10% modified ADC (2% glucose, 5% BSA, 0·85% NaCl) or in Middlebrook 7H9broth (BD Biosciences) supplemented with 10% modified ADC along with 0.05% Tween 80 without shaking.

### Determination of MIC

Minimal inhibitory concentrations (MIC) were determined using the Sensititre^™^ plates (TREK Diagnostic Systems, ThermoFisher Scientific) according to the instructions of the provider. Interpretation of MIC values followed the CLSI guidelines [10], M62.

### Whole genome sequencing

For Illumina sequencing, Paired-end (PE) DNA libraries were constructed using the Nextera XT DNA kit (Illumina, San Diego, CA, USA) according to the manufacturer’s protocol. The pooled library was prepared as recommended by the Illumina HiSeq v3 reagent preparation guide and loaded onto a cartridge (V3 chemistry) generating a 300 bp paired-end output. MinION one-dimensional (1D) libraries were constructed, using the SQK-RBK004 kit (Nanopore technologies, Oxford, UK), and loaded according to the manufacturer’s instructions onto an R9.4 flow cell. The sequencing data was collected for 48 h.

### Bioinformatic analysis

Reference genomes were *M. abscessus abscessus* (MABSa) ATCC 19977 (NC010397.1), *M. abscessus bolletii* (MABSb) CIP 108541 (NZ_JRMF00000000) and *M. abscessus massiliense* (MABSm) FLAC 047 (NZ_CP021122-1). Identification of global MABS cluster strains was achieved by including one strain of each cluster in the phylogenetic trees [BIR 948, RIVI 21, and BIR 1034 (ENA project accession ERP001039] [6]. For quality control of NGS data, the in-house pipeline QCumber (v2.1.1) (https://gitlab.com/RKIBioinformaticsPipelines/QCumber) was used. QCumber employs the software tool Trimmomatic [11], which was used for Illumina adapter removal.

All draft genomes were annotated using Prokka [12]. The determination of the maximum common genome (MCG) alignment was done by identifying the genes present in all genomes [13]. Coding sequences were clustered based on the parameters sequence similarity (min. 70%) and coverage (min. 90%) and defined the 2,085 genes that were present in each genome while fulfilling the threshold parameters as MCG. Next the allelic variants of these genes were extracted from all genomes by a BLAST-based approach, aligned individually for each gene and then concatenated, which resulted in an alignment of 2.089 Mbp. This alignment was used to calculate a maximum likelihood-based phylogeny with RAxML v.8.2.10 with 100 bootstraps under the assumption of the gtr-gamma DNA substitution model [14]. ClonalFrameML v1.11 [15] was used to correct for recombination events. The phylogenetic tree was visualized together with the distribution of accessory genes using phandango [16].

For gene variation analysis, the genome sequences of 30 MABSa isolates from one chronically infected patient collected in the years 2013 to 2017 were used to extract non-synonymous small nucleotide variants (nsSNV). The MinION sequence data together with Illumina data from an isolate originating from the first sample from this patient were used as a reference to identify nsSNVs in all other isolates compared to this genome sequence. To this end the MinION fast5 output files were demultiplexed with Deepbinner (v.0.2.0) [17]. Basecalling and barcode trimming was performed using Guppy (v.3.1.5) (Community.nanoporetech.com). The read quality was checked using pycoQC (v.2.3.1.2) [18], followed by (1) de-novo assembly of the initial sample, (2) reordering contigs against MABSa ATCC 19977 reference sequence, (3) mapping of the remaining samples from this patient against the final assembly, (4) gene annotation, (5) multi sample variant calling using the initial sample as starting point, (6) SNV and Indel filtering and (7) variant annotation. The assembly was performed using Unicycler (v0.4.7) [19] and reordered with progressiveMauve (v2.4.0) [20] against the MABSa reference sequence. Afterwards the remaining samples of the patient were mapped against the assembly using BWA (v0.7.15-r1140). Genes were annotated using prokka (v1.13.3). A multi sample variant calling was performed using GATK (v.4.1.2.0). SNVs and Indels were filtered according to GATKs best practice recommendations with few alterations. For SNVs the following filter was set: QD<2.0 || FS>60.0 || ReadPosRankSum<−8.0 || MQ<40.0 || MQRankSum<−12.5 and for indels QD<2.0 || FS>200.0 || ReadPosRankSum<−20.0. Variants that passed the filter were annotated with SnpEff (v4.3u) [21]. In a second filtering step only variants in coding regions were selected that cause non-synonymous changes ((countHom() < 30) & (DP >= 10) & ((EFF[0].IMPACT = ‘HIGH’) | (EFF[0].IMPACT = ‘MODERATE’)). Sequence analysis was supported by use of Geneious software (Geneious Prime 2020, Biomatters).

## Results

### Predominance of MABS among isolated NTM

NTM were isolated from 16 of the 42 patients (Supplementary Table S1). MABS was the most frequently isolated NTM species (11 patients) followed by *M. avium hominissuis* (MAH) (six patients), *M. intracellulare* (three patients) and *M. chimaera* (one patient). Two patients were co-infected by MABS and MAH, one patient by MABS and *M. intracellulare*, one patient by MAH and *M. intracellulare* and one patient by MABS, MAH and *M. intracellulare*.

88 isolates from the 11 patients infected with MABS were further investigated. Seven of these patients were infected with the subspecies MABSa, three with the subspecies MABSm and one patient with the subspecies MABSb. One patient had a double infection with MABSa and MABSb. 43 isolates had a rough, 43 a smooth colony morphology and two were mixed. An overview on isolates used for whole genome sequencing including subspecies, colony morphology, isolation year and accession numbers is presented in Supplementary Table S3.

### Absence of patient-to-patient transmission of MABS

Illumina NGS data were used for phylogenetic analysis of all 88 isolates from 34 respiratory samples of 11 MABS-infected patients (Supplementary Table S3). Figure 1 shows a Maximum Likelihood Tree corrected for recombination events based on 2085 core genome genes identified in the genomes of the 88 isolates and the reference strains.

**Fig. 1:**
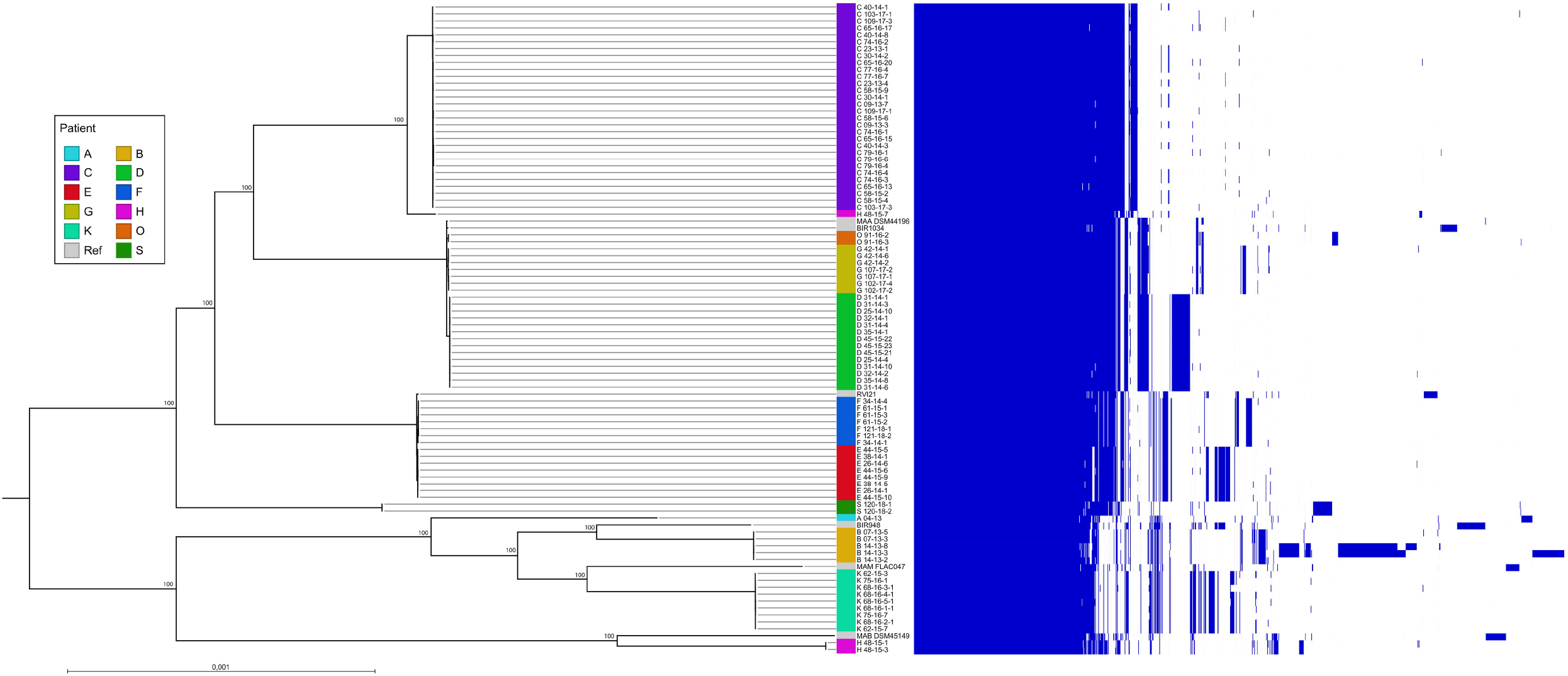
Phandango illustration showing a core genome-based Maximum likelihood tree corrected for recombination events of 88 *M. abscessus* isolates from 11 CF patients and the accessory genome. On the left side the Maximum likelihood tree is shown, on the right-side gene presence in the respective isolates is indicated by blue lines. Patients providing samples are named by letters (A to S). Names of MABS isolates are composed as follows: first letter stands for patient, first number for the sample number, second number for the year of isolation, last number for the colony number. Reference strains included in the tree were DSM 44196 (= ATCC 19977) for *M. abscessus abscessus* (accession NC_010397.1), CIP 108541 for *M. abscessus bolletii* (accession NZ_JRMF00000000.1) and FLAC047 for *M. abscessus massiliense* (accession NZ_CP021122-1). Representatives of the global patient-transmissible clusters [BIR 948 (accession ERS383065), RVI21 (accession ERS244779), BIR1034 (accession ERS383155)] described in [6] were included to identify global cluster strains present in the collection of CF isolates.

The principle tree structure was confirmed by a Maximum Likelihood Tree based on the pan genome (Supplementary Figure S1).

The phylogenetic tree shows the presence of well separated MABS clusters each belonging to one patient (A, B, C, H, K, S) as well as the presence of highly similar isolates belonging to different patients (E, F and D, G, O). One representative of each global MABS cluster [6] was included in the phylogenetic tree. While none of the patient isolates formed a cluster with the MABSm global cluster strain BIR948, isolates from patients E and F clustered with the global MABSa cluster strain RVI21, whereas isolates D, G and O clustered with global cluster strain BIR1034. Within MABSa strains up to 32 SNPs between different isolates belonging to one patient were observed in the core genome. Isolates belonging to different MABSa strains within the same global cluster exhibited SNP distances of at least 91 SNPs (cluster D/G/O/BIR1034) or at least 78 SNPs (cluster E/F/RVI21). Bryant et al. had proposed less than 20 SNPs to indicate probable recent transmission of MABS, 20-38 SNPs possible recent transmission and more than 38 SNPs no recent transmission [6]. Transmission of MABS among patients of the treatment center can thus be excluded by SNP analysis. Absence of patient-to-patient transmission was further confirmed by considerable differences in the accessory genomes (Figure 1).

### Genetic diversity evolving during persistent MABSa infection

The longest tracking time and highest sample numbers were obtained for patient C. The first respiratory sample obtained from this patient was taken in the month after diagnosis of MABS infection, followed by samples after five, 10, 18, 26, 32, 34, 36, 38, 49 and 53 months. Between two to four colonies per sample were included in the analysis. In total, 30 isolates from 11 serial sputum samples collected over 4.5 years were gained.

All 30 isolates were sequenced by Illumina technology (Supplementary Table S3, Figure 1 and Supplementary Figure S1). The Maximum Likelihood Tree in Figure 1 shows that these isolates form a cluster separated from the isolates from the other patients. One of the isolates obtained from the first sample (isolate 09-13-3) was additionally sequenced by MinION technology to complete the genome and provide a baseline reference for further analysis. Supplementary Table S4 summarizes the statistics for genome sequence assembly of the completed genome from isolate 09-13-3. The genome from isolate 09-13-3 comprised a chromosome of a size of 4.949.160 bp and a plasmid of 24.978 bp. This plasmid (pMabs 09-13, plasmid map in Supplementary Figure S2) is similar to the plasmid from strain *M. sp*. MS1601 (NCBI BlastN: 72% Query cover, 97.80 % identity).

SNV calling using the MinION genome sequence from 09-13-3 as baseline was applied to identify gene variations occurring in this patient over the 4.5-years monitoring period. Only non-synonymous (ns) SNVs and larger deletions as well as mutations in the rRNA genes were further considered. 53 genes exhibiting non-synonymous mutations with respect to isolate 09-13-3 were identified. These genes and the affected isolates are listed in Table 1.

**Table 1:**
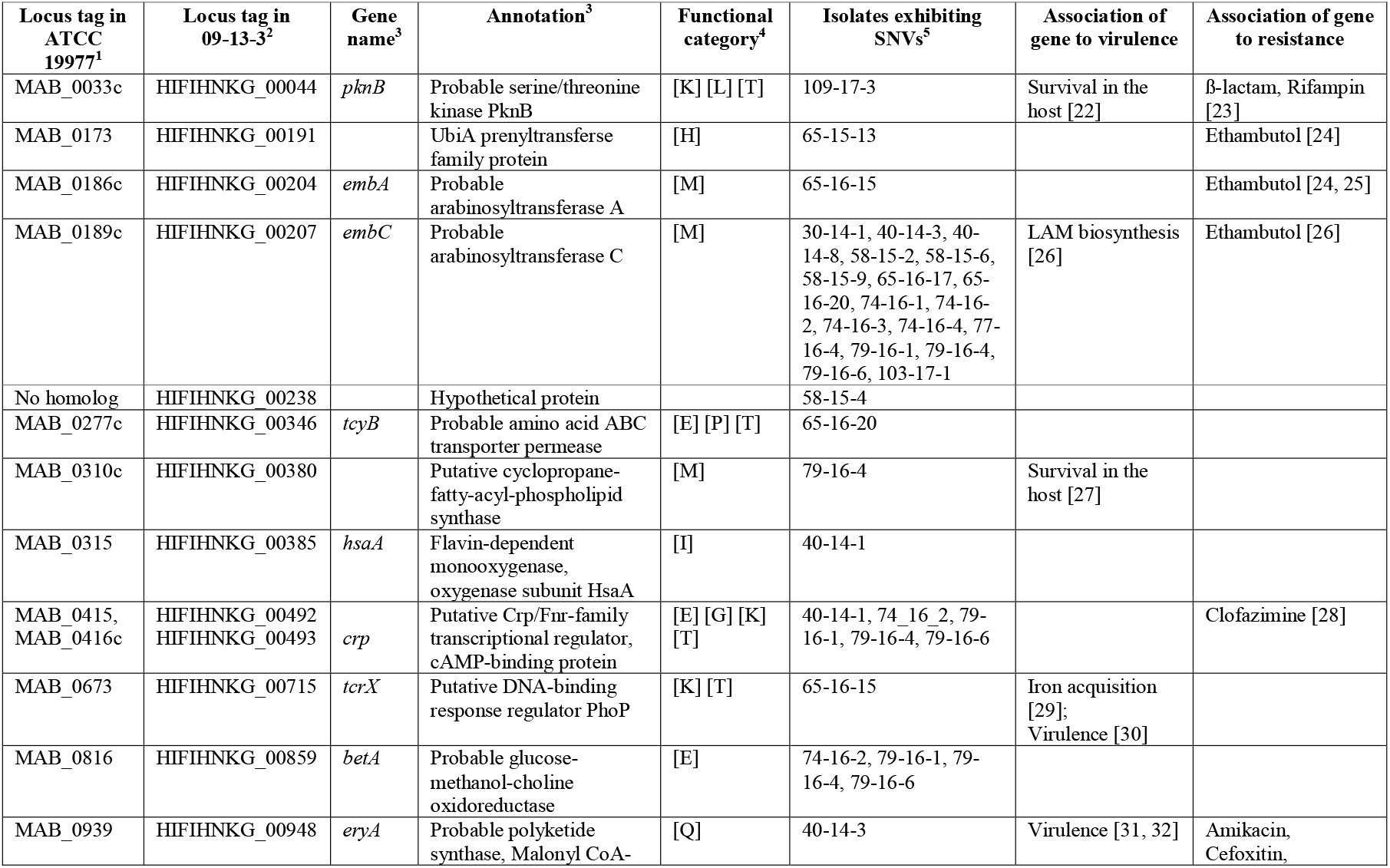
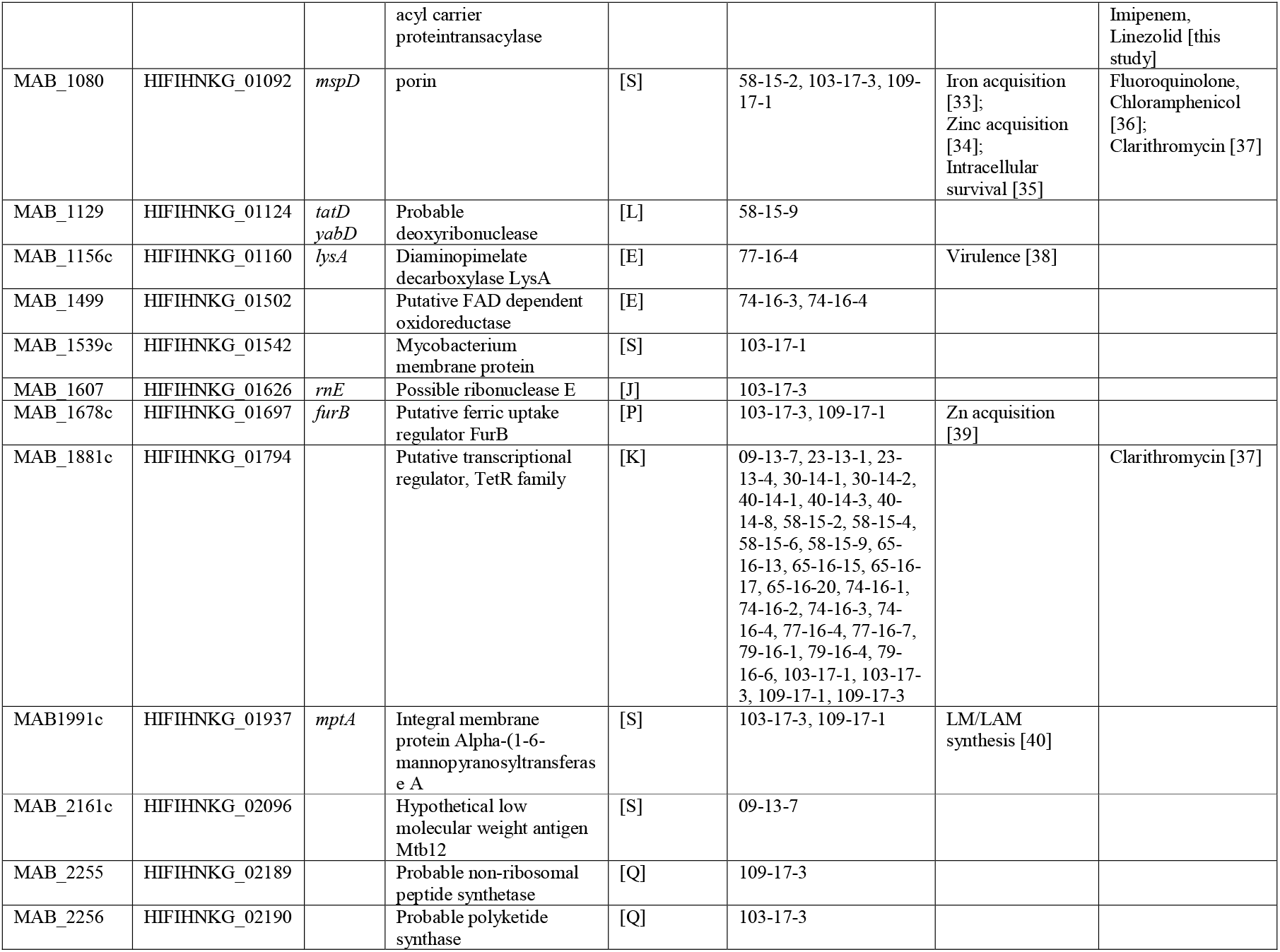
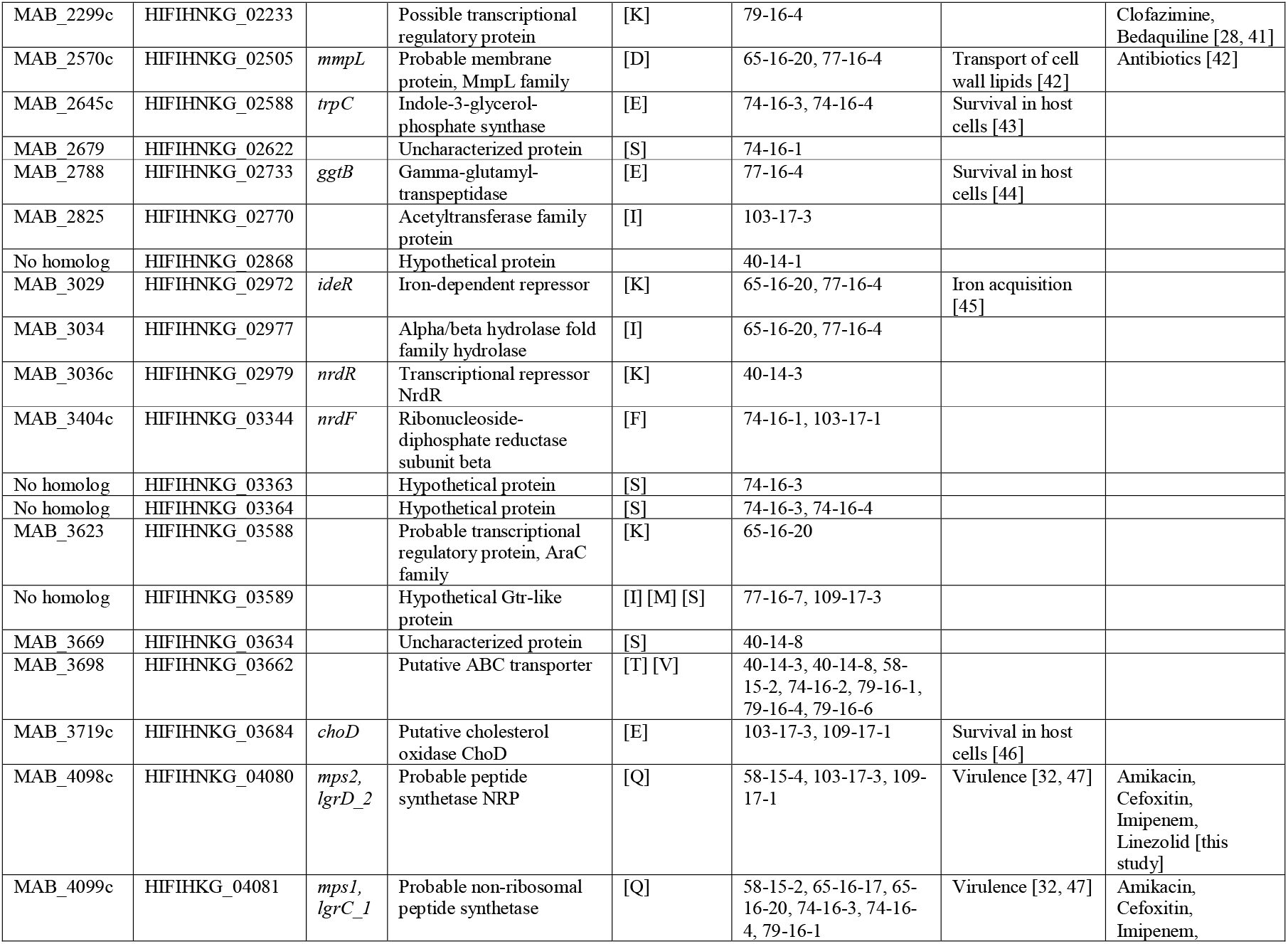
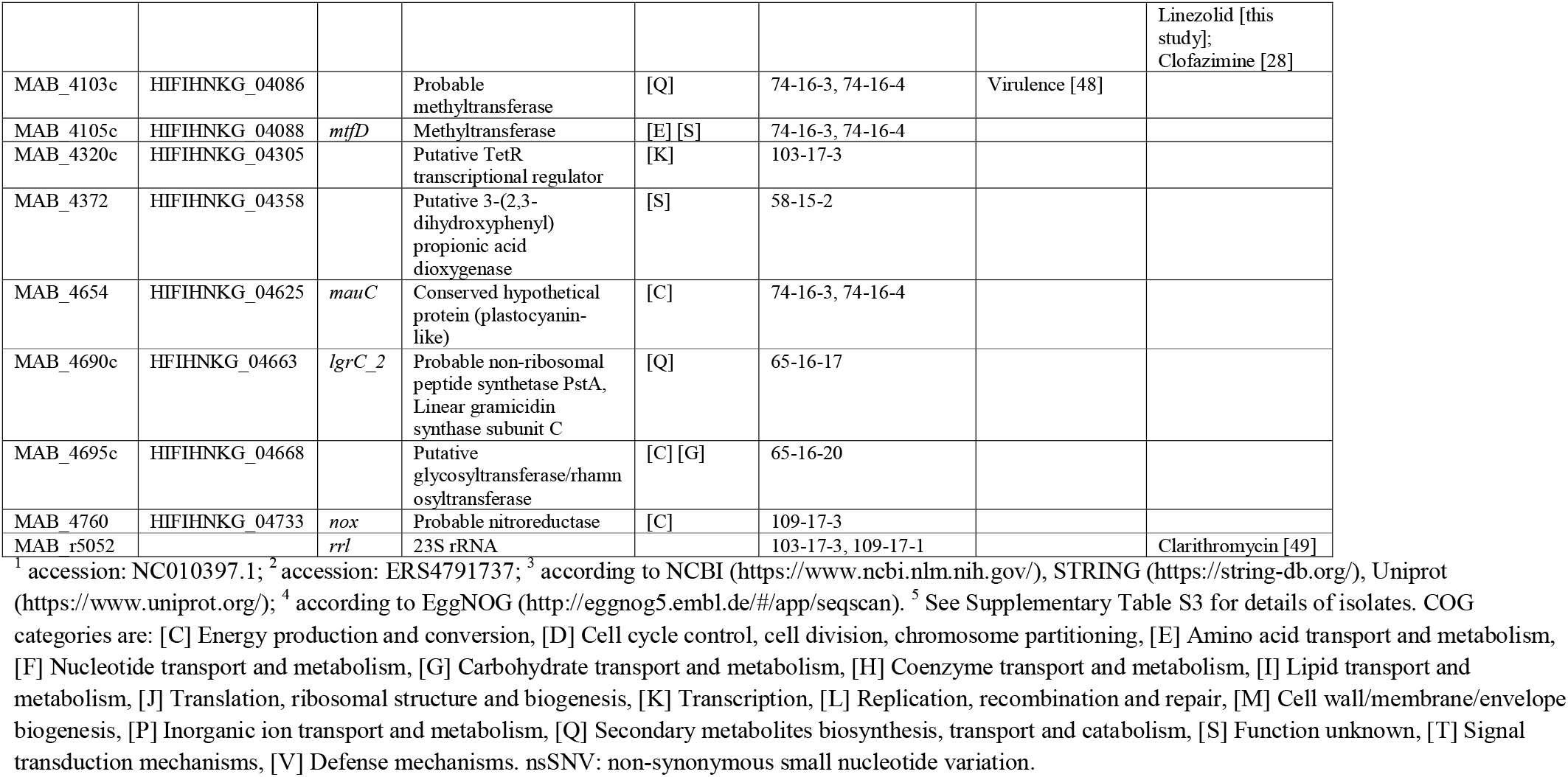
nsSNVs and larger deletions in isolates from serial samples obtained over 4.5 years from a chronically infected patient. Shown are the locus tags of the homologous genes in the reference strain ATCC 19977, the locus tags in the completed genome from the first sample (09-13-3), gene names, annotations, functional COG categories and names of the isolates exhibiting the mutations when compared to the initial isolate 09-13-3. Known associations to virulence and resistance of the genes or homologous genes in other mycobacterial species are indicated in the last two columns.

A visualization of the frequency and chronology of gene mutations occurring in the 30 isolates is provided in Figure 2. The number of mutations increased from 18 months after MABS diagnosis. Occurrence of dominating clones exhibiting and sustaining specific combinations of mutations was not observed during the observation time period. However, a clear trend was observed with respect to the loss of the plasmid pMabs over time. The strain 09-13-3 originally contained a plasmid of a size of 24.978 bp (Supplementary Figure S2). This plasmid was lost from the MABS strain in the course of chronic infection. Specifically, all isolates contained the plasmid until ten months after MABS diagnosis. However, it remained absent all in isolates sampled at and after 38 months (Figure 2).

**Figure 2:**
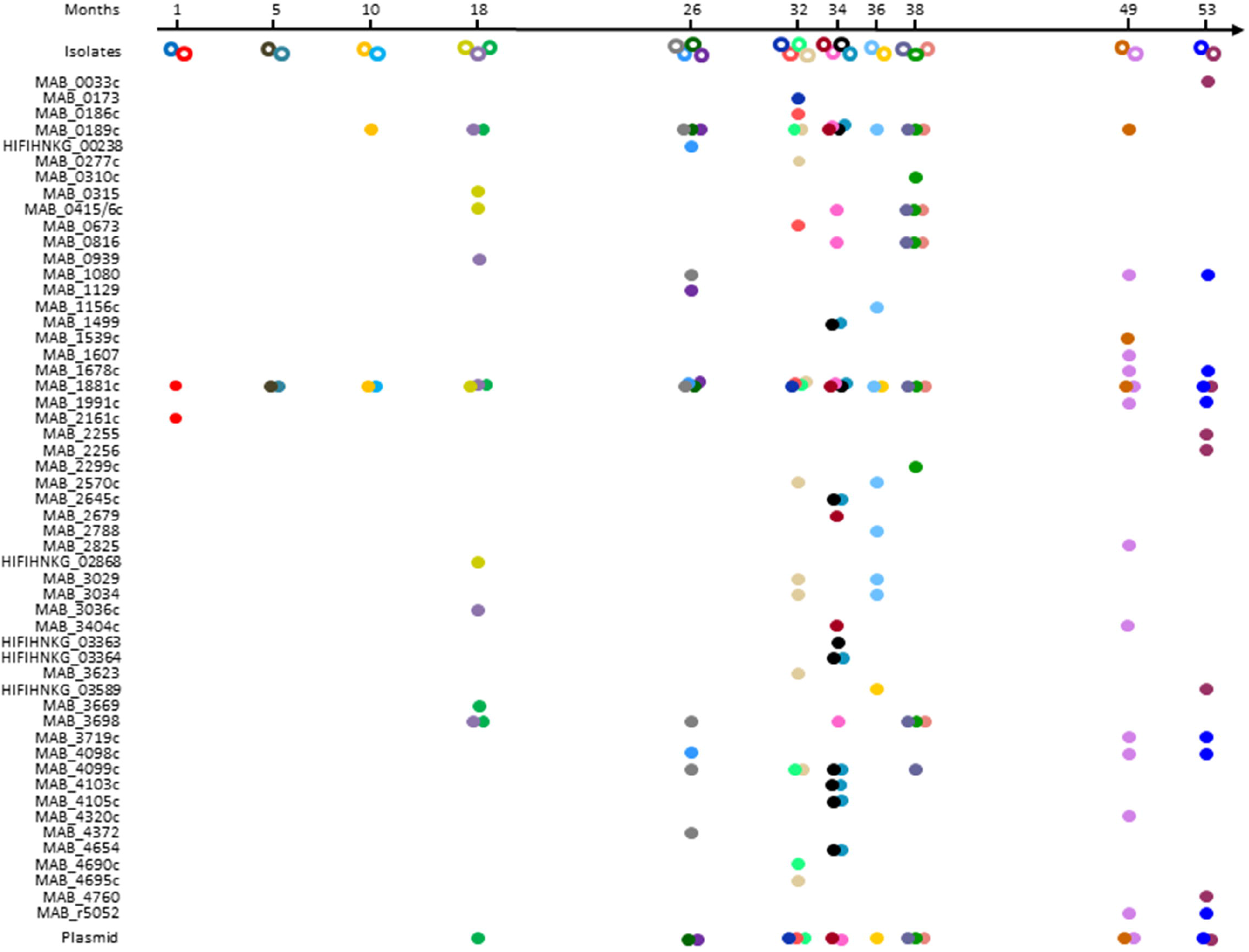
Chronology of occurrence of non-synonymous mutations in *M. abscessus* isolated during 4.5 years from patient C. The different isolates are represented as coloured open circles below the time axis. Filled circles in the lines below indicate that in this specific isolate the gene specified on the left has a non-synonymous mutation. Filled circles in the last line (Plasmid) indicate loss of the plasmid. Only in one isolate (70-16-1, 1^st^ isolate isolated after 38 months) the plasmid was not completely lost, but partly deleted.

Network analysis with the set of mutated genes by STRING (https://string-db.org/) identified ten functionally enriched PFAM protein domains (Table 2).

**Table 2:**
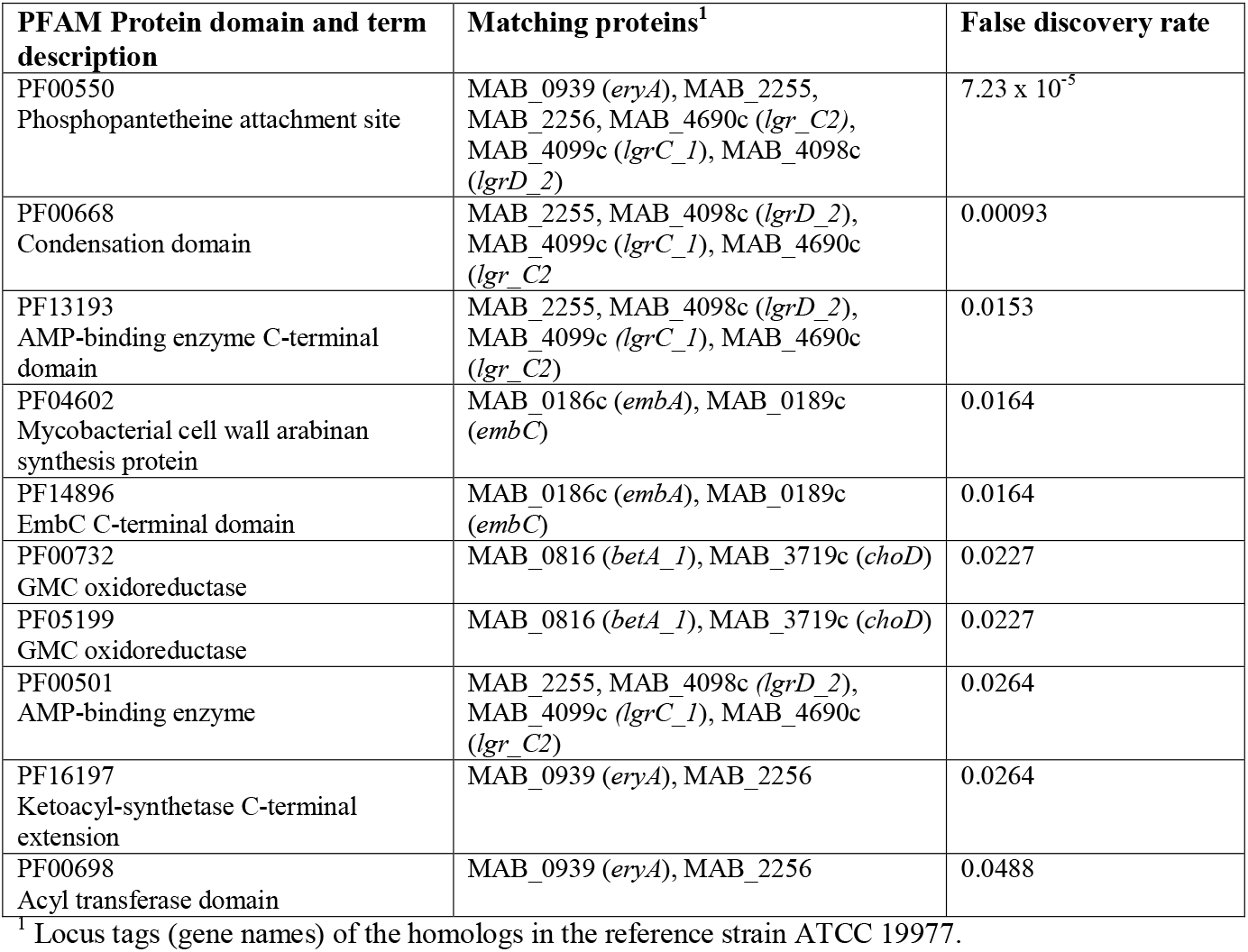
Enrichment of PFAM protein domains in the pool of genes displaying non-synonymous mutations in the *M. abscessus* isolates from patient C according to String network analysis.

Functionally enriched genes comprised a cluster involved in cell wall synthesis and resistance towards ethambutol *(embA* and *embC)* and a cluster involved in synthesis of glycopeptidolipids (GPL) *(eryA, mps2* and *mps1)*. Also, MAB_4690c belonging to a second GPL-like gene cluster from MABS [47] was among the functionally enriched proteins as well as MAB_2255 and MAB_2256, which also encode a probable non-ribosomal peptide synthetase and a probable polyketide synthase. Furthermore, *betA* (probable glucose-methanol-choline oxidoreductase) and *choD* (putative cholesterol oxidase) were among enriched genes.

The mutations in the GPL synthesis genes [50] can explain the rough morphotype of all rough isolates. Confirmation of the impact of mutations in GPL synthesis genes on GLP composition was obtained by thin layer chromatography (TLC) with GPL extracted from a smooth and four rough isolates exhibiting different types of mutations (Supplementary Figure S3).

Comparison of MICs from nine rough and nine smooth paired isolates originating from the same sputum samples showed different median MICs for Amikacin, Cefoxitin, Imipenem and Linezolid. Interestingly, rough colonies showed higher median MIC values to three of these antibiotics (Amikacin, Cefoxitin and Imipenem), while they exhibited a lower median MIC to one of them (Linezolid) (Figure 3).

**Figure 3:**
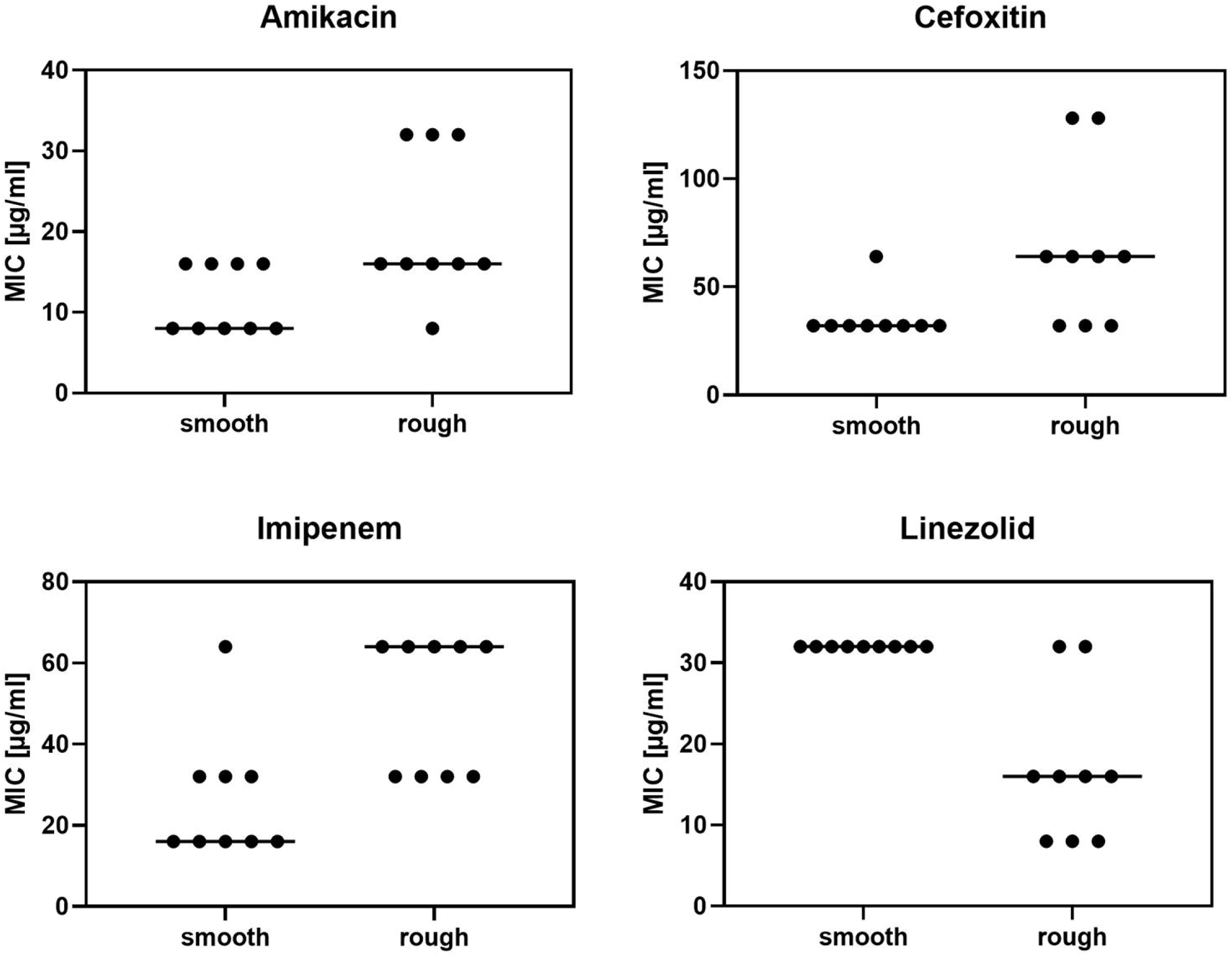
Comparison of MICs from smooth and rough isolates from patient C. Nine smooth and nine rough *M. abscessus* isolates from identical eight serial sputum samples were tested using the Sensititre system (TREK diagnostics system, ThermoFisher Scientific). Out of the 13 antibiotics available in the sensititre panel four (Amikacin, Cefoxitin, Imipenem and Linezolid) showed statistically significant differences in MIC between the smooth and rough isolates from this patient (Mann Whitney Test). Bars indicate the median values.

### Diversification in virulence- and resistance associated genes during chronic lung infection

Out of the 53 genes that had developed non-synonymous gene variations, at least 17 genes or their homologs in other mycobacteria were published to be involved in mycobacterial virulence (Table 1). These genes or their homologs exert an impact (i) on survival in the host or host cells [MAB_0033c *(pknB)*, MAB_0310c, MAB_1080 (porin), MAB_2645c *(trpC)*, MAB_2788 *(ggtB)* MAB_3719c (*choD*)], (ii) on cell wall synthesis [MAB_0189c *(embC)*, MAB_1991c *(mptA)*, MAB_2570c *(mmpL)]*, (iii) on iron and zinc acquisition [MAB_0673 *(tcrX)*, MAB_1080 (porin), MAB_1678c *(furB)*, MAB_3029 *(ideR)]*, and (iv) on GPL synthesis [MAB_0939 *(eryA)*, MAB_4098c (*mps2*), MAB_4099c (*mps1*)]. References are provided in Table 1.

Thirteen of the 53 MABS genes or their homologs in other mycobacterial species are known to be related to antibiotic resistance (Table 1). It has been shown that genes or homologs in other mycobacteria to MAB_0173, MAB_0186c *(embA)*, and 0189c *(embC)* are related to Ethambutol resistance. Clofazimine resistance was associated to genes MAB_0146c (*crp*), MAB_2299c, and MAB_4099c (*mps1*). MAB_2299c had additionally been associated with Bedaquiline resistance. Mycobacterial porin genes such as MAB_1080 were related to resistance towards, Fluoroquinolone, Chloramphenicol and Clarithromycin. Clarithromycin resistance may additionally be influenced by gene MAB_1881c and *rrl*. The mutation that was identified in the *rrl* gene (position 2270, A/C) is known to confer acquired Clarithromycin resistance. PknB from mycobacteria was shown to impact resistance towards ß-lactams and Rifampin. References are provided in Table 1.

## Discussion

MABS is a highly problematic multi-drug resistant pathogen. Despite protracted combination therapy accompanied by severe side effects, only low conversion rates of typically 40-50% are reported [8] calling for more personalized treatment also consideration of the mycobacterial population dynamics. Accordingly, the present study focused on exploring the strategy of chronic MABS to adapt to the lung environment in CF patients.

MABS predominated NTM infections in our study group, followed by MAH. The MABS subspecies distribution with 63.6% of subspecies *abscessus*, 27.3% of subspecies *massiliense* and 9% of subspecies *bolletii* was highly similar to the distribution found in the global study from Bryant et al [6].

Five of the eleven MABS-infected patients carried strains belonging to one of three global human transmissible clusters, that have been reported to be more virulent and resistant and at the same time more frequently associated with chronic disease compared to sporadic strains [6]. Nevertheless, the results from the SNP analysis ruled out transmission of MABS between patients in the study group. This was additionally supported by comparison of the accessory genomes of the isolates. Comparison of accessory genomes for discrimination of closely related MABS isolates was also proposed by Davidson [51]. Other studies (e.g. [52, 53]) had also reported the presence of global cluster strains in patient cohorts without evidence for transmission among patients. The absence of patient-to-patient transmission endorses infection control measures in the CF center in Berlin which involve among others spatial and/or temporal separation of patients with NTM in respiratory tract and a face mask wearing during the entire clinical stay [54].

Comparative genome analysis of 30 isolates from 11 serial samples from a chronically infected patient identified 53 genes with non-synonymous variations. Additionally, the plasmid from this strain present in the beginning was lost during persistent infection. This may be explained by the fitness cost of plasmid maintenance in stress conditions present in the human host and is in good accordance to a study by Shoulah et al. [55], who found more plasmid-derived genes in environmental compared to clinical isolates from MAH.

Isolates with mutations in these 53 genes are extremely valuable to study virulence and resistance mechanisms of persistent MABS, which are currently insufficiently investigated. Frequent genetic changes included those leading to GPL deficiency and rough colony morphotype. These mutations were associated with increased MICs for Amikacin, Cefoxitin and Imipenem and decreased MIC for Linezolid. In contrast to the present study comparing MICs of isogenic isolates, previous studies on strains isolated from different patients let to controversial outcomes [56, 57]. Our study emphasizes the need to give more attention to the impact of morphotypes on drug resistance when searching for new anti-mycobacterial drugs.

Of the 53 genes exhibiting genetic diversity, at least 23 genes or their homologs in other mycobacteria have been assigned to be virulence- and/or resistance-associated. Genetic diversity evolved in genes related to resistance to Ethambutol, Rifampin, Clofazimine, Bedaquiline, Fluoroquinolone, Chloramphenicol, Imipenem, Cefoxitin and Clarithromycin. Two isolates had acquired the A to C mutation at position 2270 (MABS numbering) in the *rrl* gene, a mutation known to confer acquired Clarithromycin resistance [58].

Interestingly, in-patient evolution did not bring forth fewer dominating sub-populations but rather fostered the co-existence of diverse mutant sub-populations. The upsurge of genetic diversity possibly enables the population to adapt to changing living conditions as illustrated by the following example.

MAB ecology combines the ability to survive both in the environment and in human airways, which offer disparate access to biometals such as zinc and iron. Such metals are, on the one hand, essential co-factors of enzymes and structural components of regulatory proteins. On the other hand, however, excessive concentrations thereof can be toxic. Therefore, a stringent regulatory system for homeostasis is required. The sputum from CF patients displays highly enriched metal concentrations of zinc and iron [59], which may favor genetic diversity within genes involved in zinc and iron homeostasis during chronic infection. Zinc uptake in mycobacteria is regulated by *zur-smtB*. Among the genes exhibiting gene diversity in persistent infection, *zur* is a zinc-binding repressor controlling genes involved in zinc uptake [34]. The porin MspD was shown to be induced by zinc starvation or *zur* deletion [34]. Moreover, MAB_1080, which was annotated as the porin protein MspD, was also among the genes exhibiting diversity upon chronic infection. Mutations were also identified in *ideR*, a gene that regulates transcription in response to iron levels [60], as well as in *tcrX*, which was shown to be up-regulated when *M. tuberculosis* was grown under iron-limited conditions [29]. Furthermore, Msp porins of rapid growing mycobacteria promote growth in nutrient-limited conditions by enhancing diffusion of small hydrophilic molecules into the cells. At the same time, however, they limit intracellular survival by increasing vulnerability to killing by reactive nitrogen [35, 61]. Therefore, the mutation in MAB_1080 may additionally impact the survival of MABS in macrophages.

ChoD, a cholesterol oxidase, is needed in *M. tuberculosis* survival in macrophages [46]. A mutant deficient in the gamma-glutamyl-transpeptidase (GgtB) from *M. tuberculosis*, which is homologous to MAB_2788, was shown to be resistant to the toxic effects of Glutathione/ S-nitrosoglutathione and therefore better survived in macrophages [44]. Macrophages are not the only host cells for mycobacteria, some of which are also able to replicate in Type II alveolar epithelial cells (AECs). The tryptophan synthesis gene *trpC* from *M. tuberculosis* is strongly up-regulated during growth in AECs [43] and also the homologue from this gene was mutated in two of the MABS isolates.

MptA, EmbC and MmpL family proteins are involved in lipid/glycolipid synthesis. MptA is a mannosyltransferase necessary for synthesis of the mannan backbone from Lipomannan. EmbC from *M. tuberculosis* catalyses arabinosylation of Lipomannan to form Lipoarabinomannan (LAM), which is involved in immune response by interacting with TLR2 and mannose-receptor [26, 40, 62]. Interestingly, a longitudinal analysis by Kreutzfeld [63] of 6 MABS *bolletii* isolates from a CF patient collected over 11 years and also identified mutations in the *embABC* operon during chronic infection.

In conclusion, our data indicate that the survival strategy of MABS in the CF lung is not towards the clonal expansion of few dominant variants but the preservation of heterogeneous subpopulations allowing adaptation to changing lung conditions. Similar future longitudinal studies involving other MABS strains and patients will further explore the range of variation of MABS in-patient evolution during chronic lung infection. Our study suggests that intervention procedures against MABS should target MABS populations instead of specific isolates.

## Acknowledgements

We would like to thank the patients with CF, their families and the team of the CF center in Berlin. We also thank Andrea Thürmer (Robert Koch Institute Berlin) and her team for genome sequencing and Nils Höpner (Robert Koch Institute Berlin) for experimental support. The work from Carsten Schwarz was supported by a German Infectiology Award.

## Funding

The work from Carsten Schwarz was supported by a German Infectiology Award.

## Disclosure of interest

The authors report no conflict of interest.

## Data Availability Statement

All genome sequences have been submitted to the European Nucleotide Archive (ENA).

## Data deposition

Genome sequences are deposited at the European Nucleotide Archive (ENA).

## Ethical statement

Permission for the study was obtained from the ethics committee of the Charité –Universitätsmedizin Berlin (EA2/093/12). Written consent had been obtained from all patients.

## Statement to health and safety regulations

The authors confirm compliance with national health and safety regulations.

## Legends for Supplementary Figures

Supplementary Figure S1:

Pan-genome-based maximum likelihood tree of 88 *M. abscessus* isolates from 11 CF patients and reference strains. Patients are named by letters (A to S). Names of *M. abscessus* isolates are composed as follows: first letter stands for patient, first number for the year of isolation, last number for the colony number. Reference strains included in the tree were DSM 44196 (= ATCC 19977) for *M. abscessus abscessus* (accession NC_010397.1), CIP 108541 for *M. abscessus bolletii* (accession NZ_JRMF00000000) and FLAC047 for *M. abscessus massiliense* (accession NZ_CP021122-1). Representatives of the global patient-transmissible clusters [BIR 948 (accession ERS383065), RVI21 (accession ERS244779), BIR1034 (accession ERS383155)] described by [1] were included to identify global cluster strains present in the collection of CF isolates. The scale of the bar represents the percentage of substitutions per site.

Supplementary Fig. S2:

Plasmid map with annotations of plasmid pMabs-09-13 from isolate 09-13-3.

Supplementary Fig. S3:

Thin layer chromatography (TLC) of extracted glycopeptidolipids (GPL) from smooth and rough *M. abscessus* isolates. GPL were extracted from strains 58-15-4 (A), 40-14-3 (B), 74-16-3 (C) and 65-16-17 (D) and subjected to TLC as described in Fujiwara et al. [PLoS One 2015, 10(5):e0126813]. Isolate 58-15-4 had a mutation *mps2*, isolate 40-14-3 in *eryA*, isolate 74-16-3 in *mps1, mtfD* and MAB_4103c and isolate 65-16-17 in *mps1* and MAB_4690c (Table 2). The different mutants displaying a rough phenotype showed deviating GPL patterns when compared to the wildtype (E, isolate 23-13-1).

## Captions of Supplementary Tables

Supplementary Table S1:

Patient characteristics and NTM isolation.

Supplementary Table S2:

Primers used for identification of NTM.

Supplementary Table S3:

List of isolates used for whole genome sequencing with accession numbers.

Supplementary Table S4:

Statistics for genome sequence assembly for isolate MABS 09-13-3 (MinION, accession ERS4791737).

## Notes

https://www.ebi.ac.uk/ena/browser/view/PRJEB39129

### Competing Interest Statement

The authors have declared no competing interest.

