## Supplementary Material for "Genetic diversification of persistent *Mycobacterium abscessus* within Cystic Fibrosis patients"

**Supplementary Fig. S1**

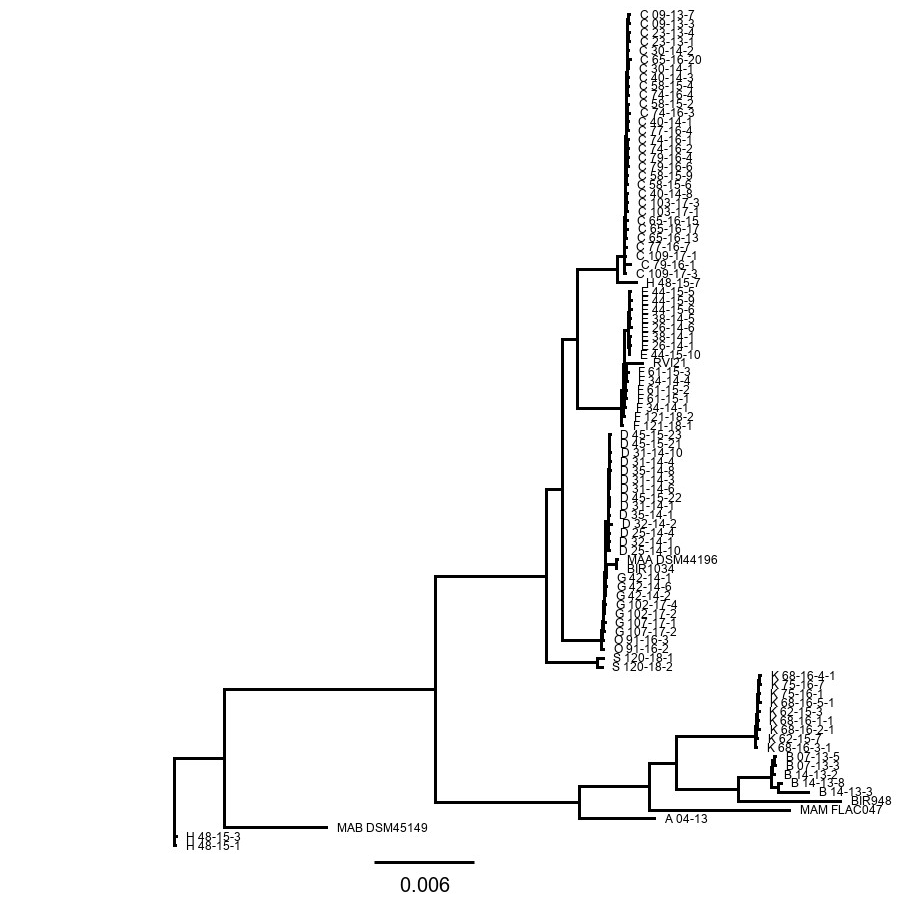

Pan-genome-based maximum likelihood tree of 88 *M. abscessus* isolates from 11 CF patients and reference strains. Patients are named by letters (A to S). Names of *M. abscessus* isolates are composed as follows: first letter stands for patient, first number for the year of isolation, last number for the colony number. Reference strains included in the tree were DSM 44196 (= ATCC 19977) for *M. abscessus abscessus* (accession NC_010397.1), CIP 108541 for *M. abscessus* *bolletii* (accession NZ_JRMF00000000) and FLAC047 for *M. abscessus* *massiliense* (accession NZ_CP021122-1). Representatives of the global clusters [BIR 948 (accession ERS383065), RVI21 (accession ERS244779), BIR1034 (accession ERS383155)] described by [1] were included to identify global cluster strains present in the collection of CF isolates. The scale of the bar represents the percentage of substitutions per site.

**Method**
A pan genome of the three reference genomes was constructed using Seq-Seq-Pan [2]. In the next step, the adapter and quality trimmed samples were mapped against the pan genome using BWA (v0.7.15-r1140) (https://arxiv.org/abs/1303.3997). Variant calling was performed with GATK (v4.1.2.0) [3] according to GATKs best practice pipeline with default parameters. The variants were used to generate consensus sequences of each sample with BCFtools (v1.3.2) https://doi.org/10.1093/bioinformatics/btx100). A multiple sequence alignment was performed with Seq-Seq-Pan [2] using the consensus sequences of the samples, the three cluster references and the three references of *M. abscessus* used for the pan genome. In the final step the phylogeny of the samples was inferred based on the alignment using PhyML (v3.3.20190909) [4]. The HYK85 substitution model was applied and 100 bootstrap replicates were used to evaluate the confidence in nodes.

1.	Bryant JM, Grogono DM, Rodriguez-Rincon D, Everall I, Brown KP, Moreno P, Verma D, Hill E, Drijkoningen J, Gilligan P et al: Emergence and spread of a human-transmissible multidrug-resistant nontuberculous mycobacterium. Science 2016, 354(6313):751-757.

2.	Jandrasits C, Dabrowski PW, Fuchs S, Renard BY: seq-seq-pan: building a computational pan-genome data structure on whole genome alignment. BMC Genomics 2018, 19(1):47.

3.	McKenna A, Hanna M, Banks E, Sivachenko A, Cibulskis K, Kernytsky A, Garimella K, Altshuler D, Gabriel S, Daly M et al: The Genome Analysis Toolkit: a MapReduce framework for analyzing next-generation DNA sequencing data. Genome Res 2010, 20(9):1297-1303.

4.	Guindon S, Dufayard JF, Lefort V, Anisimova M, Hordijk W, Gascuel O: New algorithms and methods to estimate maximum-likelihood phylogenies: assessing the performance of PhyML 3.0. Syst Biol 2010, 59(3):307-321.

**Supplementary Figure S2**

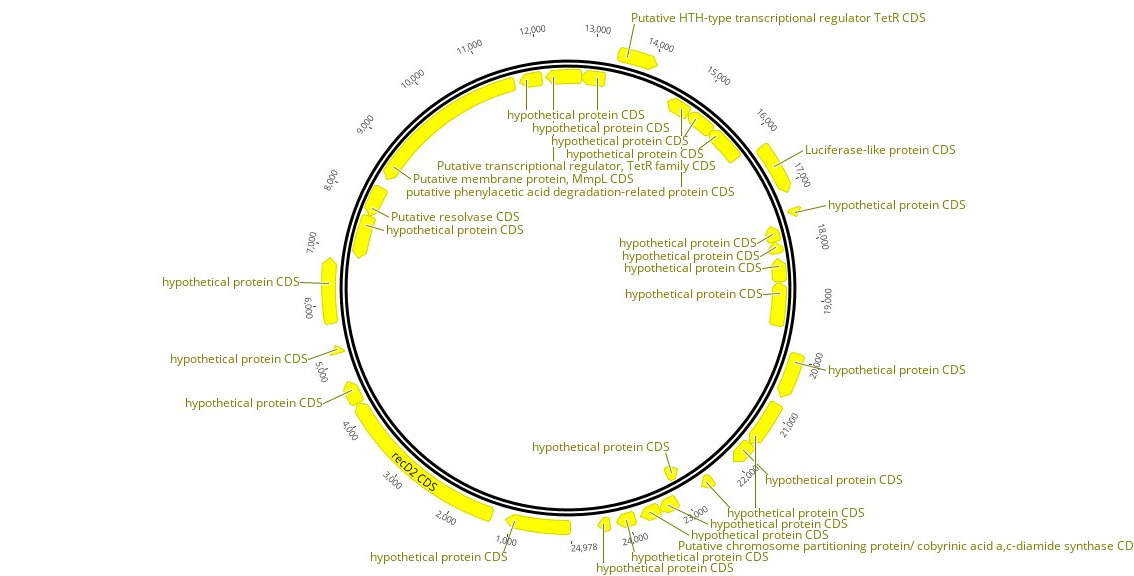

Plasmid map with annotations of plasmid pMabs-09-13 from isolate 09-13-3.

**Supplementary Figure S3**

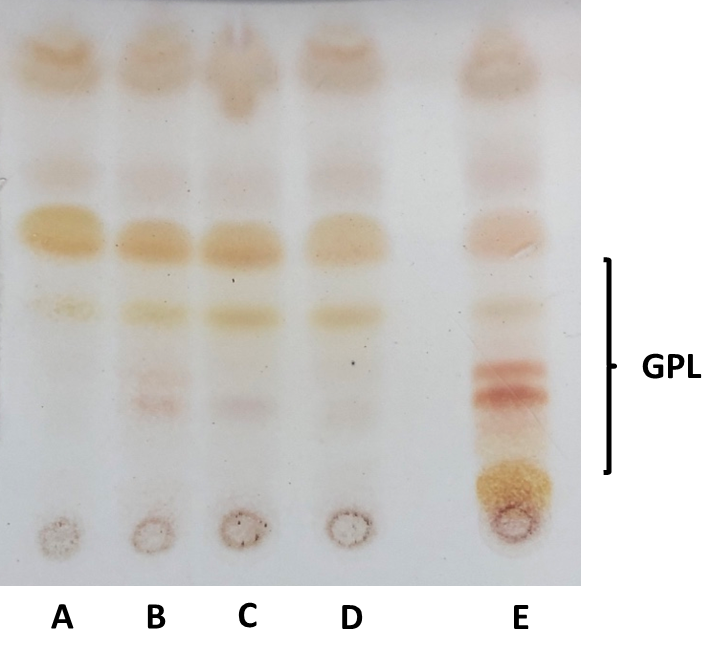

Thin layer chromatography (TLC) of extracted Glycopeptidolipids (GPL) from smooth and rough *M. abscessus* isolates. GPL were extracted from strains 58-15-4 (A), 40-14-3 (B), 74-16-3 (C) and 65-16-17 (D) and subjected to TLC as described in Fujiwara et al. [PLoS One 2015, 10(5):e0126813]. Isolate 58-15-4 had a mutation *mps2*, isolate 40-14-3 in *eryA*, isolate 74-16-3 in *mps1*, *mtfD* and MAB_4103c and isolate 65-16-17 in *mps1* and MAB_4690c (Table 2). The different mutants displaying a rough phenotype showed deviating GPL patterns when compared to the wildtype (E, isolate 23-13-1).

**Supplementary Table S1:** Patient characteristics and NTM isolation.

|  | **All patients** | **Without NTM** | **With NTM** |
| --- | --- | --- | --- |
| All | 42 | 26 | 16 |
| Male | 20 | 9 | 11 |
| Female | 22 | 17 | 5 |
| Median age (range)^1^ | 26 (1-51) | 26 (1-51) | 28 (8-48) |
| Median BMI (range)^1^ | 20.1 (15.2-26.4) | 20.0 (16.8-23.2) | 20.9 (15.2-26.4) |
| Median FEV1 [%] (range)^1,2^ | 55.0 (21-110) | 52.5 (21-110) | 62 (25-106) |
| Pancreas insufficiency^1^ | 41 | 25 | 16 |
| Diabetes^1^ | 18 | 12 | 6 |
| *Pseudomonas aeruginosa* | 19 | 13 | 6 |
| Organ transplantation | 5 | 4 | 1 |

^1^ Data refer to the first sampling if serial samples were taken. ^2^ % of predicted.

**Supplementary Table S2:** Primers used for identification of NTM.

| Primer | sequence | Target Gene and Organism | Anneal. Temp. [°C] | Product size [bp] | Reference |
| --- | --- | --- | --- | --- | --- |
| 16SrRNA-gene_F | ATAAGCCTGGGAAACTGGGT | 16rRNA Mycobacteria | 60 | 484 | [1] |
| 16SrRNA-gene_R | CACGCTCACAGTTAAGCCGT |  |  |  |  |
| 16SrRNA-gene_2_F | TGGAGAGTTTGATCCTGGCTCAG | 16rRNA  Mycobacteria | 57.5 | 1030 | [2] |
| 16SrRNA-gene_2_R | TGCACACAGGCCACAAGGGA |  |  |  |  |
| 16S-1511f  23S-23S | AAGTCGTAACAAGGTAGCCG  TCGCCAAGGCATCCACC | ITS Mycobacteria | 53 | 389 | [3] |
| dnaJ-1 | GACTTCTACAAGGAGCTGGG | *dnaJ M. avium, M. intracellulare* | 58 | 140 | [4] |
| dnaJ-2 | GAGACCGCCTTGAATCGTTC |  |  |  |  |
| IS1245-1 | GAGTTGACCGCGTTCATCG | *M. avium avium M. avium silvaticum M. avium hominissuis* | 58 | 385 | [4] |
| IS1245-2 | CGTCGAGGAAGACATACGG |  |  |  |  |
| IS901-1 | GGATTGCTAACCACGTGGTG | IS901 *M. avium avium*  *M. avium silvaticum* | 58 | 577 | [4] |
| IS901-2 | GCGAGTTGCTTGATGAGCG |  |  |  |  |
| Mab1 for | CCTCGAGCCCAAGATCTGTC | VNTR11 *M. abscessus* | 63 | 393-492 | [5] |
| Mab1 rev | ATACCGGGATACGCCAAGAT |  |  |  |  |
| Mab2 for | AAGGGACTGGGACTGATCG | VNTR23 *M.abscessus* | 63 | 196-238 | [5] |
| Mab2 rev | CCGGAGACCGACCTCTTC |  |  |  |  |
| Erm_FW Erm_BW | GACCGGGGCCTTCTTCGTGAT  GACTTCCCCGCACCGATTCC | *erm*(41) *M.abscessus complex* | 62/66 | ~700/ ~350 | [6] |

1.	Shin SJ, Lee BS, Koh WJ, Manning EJB, Anklam K, Sreevatsan S, Lambrecht RS, Collins MT: Efficient differentiation of Mycobacterium avium complex species and subspecies by use of five-target multiplex PCR. J Clin Microbiol 2010, 48(11):4057-4062.

2.	Rogall T, Flohr T, Böttger EC: Differentiation of Mycobacterium species by direct sequencing of amplified DNA. Journal of general microbiology 1990, 136(9):1915-1920.

3.	Harmsen D, Dostal S, Roth A, Niemann S, Rothgänger J, Sammeth M, Albert J, Frosch M, Richter E: RIDOM: Comprehensive and public sequence database for identification of Mycobacterium species. BMC Infect Dis 2003, 3.

4.	Moravkova M, Hlozek P, Beran V, Pavlik I, Preziuso S, Cuteri V, Bartos M: Strategy for the detection and differentiation of Mycobacterium avium species in isolates and heavily infected tissues. Res Vet Sci 2008, 85(2):257-264.

5.	Choi GE, Chang CL, Whang J, Kim HJ, Kwon OJ, Koh WJ, Shin SJ: Efficient differentiation of Mycobacterium abscessus complex isolates to the species level by a novel PCR-based variable-number tandem-repeat assay. J Clin Microbiol 2011, 49(3):1107-1109.

6.	Shallom SJ, Gardina PJ, Myers TG, Sebastian Y, Conville P, Calhoun LB, Tettelin H, Olivier KN, Uzel G, Sampaio EP et al: New rapid scheme for distinguishing the subspecies of the Mycobacterium abscessus group and identifying Mycobacterium massiliense isolates with inducible clarithromycin resistance. J Clin Microbiol 2013, 51(9):2943-2949.

**Supplementary Table S3:** List of isolates used for whole genome sequencing with accession numbers.

| Isolate name | Patient | Year of isolation | Subspecies | Colony morphology | Accession |
| --- | --- | --- | --- | --- | --- |
| 04/13 | A | 2013 | massiliense | rough | ERS4791649 |
| 07-13-3 | B | 2013 | massiliense | rough | ERS4791650 |
| 07-13-5 | B | 2013 | massiliense | rough | ERS4791651 |
| 14-13-2 | B | 2013 | massiliense | rough | ERS4791654 |
| 14-13-3 | B | 2013 | massiliense | rough | ERS4791655 |
| 14-13-8 | B | 2013 | massiliense | rough | ERS4791656 |
| 09-13-3 | C | 2013 | abscessus | smooth | ERS4791737 (MinION)  ERS4791736 (Illumina) ERS4791652 (Illumina) |
| 09-13-7 | C | 2013 | abscessus | smooth | ERS4791653 |
| 23-13-1 | C | 2013 | abscessus | smooth | ERS4791657 |
| 23-13-4 | C | 2013 | abscessus | smooth | ERS4791658 |
| 30-14-1 | C | 2014 | abscessus | smooth | ERS4791663 |
| 30-14-2 | C | 2014 | abscessus | smooth | ERS4791664 |
| 40-14-1 | C | 2014 | abscessus | smooth | ERS4791679 |
| 40-14-3 | C | 2014 | abscessus | rough | ERS4791677 |
| 40-14-8 | C | 2014 | abscessus | smooth | ERS4791678 |
| 58-15-2 | C | 2015 | abscessus | rough | ERS4791693 |
| 58-15-4 | C | 2015 | abscessus | rough | ERS4791694 |
| 58-15-6 | C | 2015 | abscessus | smooth | ERS4791695 |
| 58-15-9 | C | 2015 | abscessus | smooth | ERS4791696 |
| 65-16-13 | C | 2016 | abscessus | smooth | ERS4791702 |
| 65-16-15 | C | 2016 | abscessus | smooth | ERS4791703 |
| 65-16-17 | C | 2016 | abscessus | rough | ERS4791704 |
| 65-16-20 | C | 2016 | abscessus | rough | ERS4791705 |
| 74-16-1 | C | 2016 | abscessus | smooth | ERS4791713 |
| 74-16-2 | C | 2016 | abscessus | smooth | ERS4791714 |
| 74-16-3 | C | 2016 | abscessus | rough | ERS4791715 |
| 74-16-4 | C | 2016 | abscessus | rough | ERS4791716 |
| 77-16-4 | C | 2016 | abscessus | smooth | ERS4791717 |
| 77-16-7 | C | 2016 | abscessus | smooth | ERS4791718 |
| 79-16-1 | C | 2016 | abscessus | rough | ERS4791719 |
| 79-16-4 | C | 2016 | abscessus | smooth | ERS4791720 |
| 79-16-6 | C | 2016 | abscessus | smooth | ERS4791721 |
| 103-17-1 | C | 2017 | abscessus | smooth | ERS4791727 |
| 103-17-3 | C | 2017 | abscessus | rough | ERS4791726 |
| 109-17-1 | C | 2017 | abscessus | rough | ERS4791730 |
| 109-17-3 | C | 2017 | abscessus | smooth | ERS4791731 |
| 25-14-4 | D | 2014 | abscessus | smooth | ERS4791659 |
| 25-14-10 | D | 2014 | abscessus | smooth | ERS4791660 |
| 31-14-1 | D | 2014 | abscessus | smooth | ERS4791665 |
| 31-14-3 | D | 2014 | abscessus | smooth | ERS4791666 |
| 31-14-4 | D | 2014 | abscessus | smooth | ERS4791667 |
| 31-14-6 | D | 2014 | abscessus | smooth | ERS4791668 |
| 31-14-10 | D | 2014 | abscessus | smooth | ERS4791669 |
| 32-14-1 | D | 2014 | abscessus | rough | ERS4791670 |
| 32-14-2 | D | 2014 | abscessus | rough | ERS4791671 |
| 35-14-1 | D | 2014 | abscessus | rough | ERS4791673 |
| 35-14-8 | D | 2014 | abscessus | rough | ERS4791674 |
| 45-15-21 | D | 2015 | abscessus | rough | ERS4791687 |
| 45-15-22 | D | 2015 | abscessus | rough | ERS4791688 |
| 45-15-23 | D | 2015 | abscessus | rough | ERS4791689 |
| 26-14-1 | E | 2014 | abscessus | rough | ERS4791661 |
| 26-14-6 | E | 2014 | abscessus | rough | ERS4791662 |
| 38-14-1 | E | 2014 | abscessus | rough | ERS4791675 |
| 38-14-5 | E | 2014 | abscessus | rough | ERS4791676 |
| 44-15-5 | E | 2015 | abscessus | rough | ERS4791683 |
| 44-15-6 | E | 2015 | abscessus | rough | ERS4791684 |
| 44-15-9 | E | 2015 | abscessus | rough | ERS4791685 |
| 44-15-10 | E | 2015 | abscessus | rough | ERS4791686 |
| 34-14-1 | F | 2014 | abscessus | rough | RRCG00000000 |
| 34-14-4 | F | 2014 | abscessus | rough | ERS4791672 |
| 61-15-1 | F | 2015 | abscessus | rough | ERS4791697 |
| 61-15-2 | F | 2015 | abscessus | rough | ERS4791698 |
| 61-15-3 | F | 2015 | abscessus | rough | ERS4791699 |
| 121-18-1 | F | 2018 | abscessus | rough | ERS4791734 |
| 121-18-2 | F | 2018 | abscessus | rough | ERS4791735 |
| 42-14-1 | G | 2014 | abscessus | rough | ERS4791680 |
| 42-14-2 | G | 2014 | abscessus | smooth | ERS4791681 |
| 42-14-6 | G | 2014 | abscessus | smooth | ERS4791682 |
| 102-17-2 | G | 2017 | abscessus | rough | ERS4791724 |
| 102-17-4 | G | 2017 | abscessus | rough | ERS4791725 |
| 107-17-1 | G | 2017 | abscessus | rough | ERS4791728 |
| 107-17-2 | G | 2017 | abscessus | rough | ERS4791729 |
| 48-15-1 | H | 2015 | bolletii | smooth | ERS4791690 |
| 48-15-3 | H | 2015 | bolletii | smooth | ERS4791691 |
| 48-15-7 | H | 2015 | abscessus | smooth | ERS4791692 |
| 62-15-3 | K | 2015 | massiliense | smooth | ERS4791701 |
| 62-15-7 | K | 2015 | massiliense | smooth | ERS4791700 |
| 68-16-1-1 | K | 2016 | massiliense | smooth | ERS4791706 |
| 68-16-2-1 | K | 2016 | massiliense | smooth | ERS4791707 |
| 68-16-3-1 | K | 2016 | massiliense | smooth | ERS4791708 |
| 68-16-4-1 | K | 2016 | massiliense | smooth | ERS4791709 |
| 68-16-5-1 | K | 2016 | massiliense | smooth | ERS4791710 |
| 75-16-1 | K | 2016 | massiliense | mix | ERS4791711 |
| 75-16-7 | K | 2016 | massiliense | mix | ERS4791712 |
| 91-16-2 | O | 2016 | abscessus | rough | ERS4791722 |
| 91-16-3 | O | 2016 | abscessus | rough | ERS4791723 |
| 120-18-1 | S | 2018 | abscessus | smooth | ERS4791732 |
| 120-18-2 | S | 2018 | abscessus | smooth | ERS4791733 |

**Supplementary Table S4:** Statistics for genome sequence assembly for isolate MABS 09-13-3 (MinION, accession ERS4791737).

| **Statistics without reference** |  |
| --- | --- |
| N50 | 4949160 bp |
| N75 | 4949160 bp |
| L50 | 1 |
| L75 | 1 |
| Number of contigs | 2  (Chromosome: 4.949.160 bp Plasmid: 24.978 bp) |
| GC [%] | 64.27 |
| **Statistics with reference *M. abscessus abscessus* NC010397_1** |  |
| Genome fraction of reference covered [%] | 93.124 |
| Largest alignment | 572753 bp |
| Total aligned length | 4717931 bp |

Quast (v 5.0.0) [1] was used for assembly quality control.

[1] Mikheenko A, Prjibelski A, Saveliev V, Antipov D, Gurevich A: Versatile genome assembly evaluation with QUAST-LG. Bioinformatics (2018) 34 (13): i142-i150. doi: 10.1093/bioinformatics/bty266.
